# An orphan viral genome with controversial evolutionary status sheds light on a distinct lineage of flavi-like virus infecting plants

**DOI:** 10.1101/2024.05.27.596083

**Authors:** Zhongtian Xu, Luping Zheng, Fangluan Gao, Yiyuan Li, Zongtao Sun, Jianping Chen, Chuanxi Zhang, Junmin Li, Xifeng Wang

**Author notes:** Corresponding author (J. Li), (X. Wang).

## Abstract

Advancements in high-throughput sequencing and associated bioinformatics methods have significantly expanded the RNA virus repertoire, including novel viruses with highly divergent genomes encoding ‘orphan’ proteins that apparently lack homologous sequences. This absence of homologs in routine sequence similarity search complicates their taxonomic classification and raises a fundamental question: Do these orphan viral genomes represent *bona fide* viruses? In 2022, an orphan viral genome encoding a large polyprotein was identified in alfalfa (*Medicago sativa*) and named Snake River alfalfa virus (SRAV). Initially, SRAV was proposed to be within the flavi-like lineage of the family *Flaviviridae*. Subsequently, another research group showed its common occurrence in alfalfa but challenged its taxonomic position, suggesting it belongs to the family *Endornaviridae* rather than *Flaviviridae*. In this study, a large-scale analysis of 77 publicly available small RNA datasets indicated that SRAV could be detected across various tissues and cultivars of alfalfa, and has a broad geographical distribution. Moreover, profiles of the SRAV-derived small interfering RNAs (vsiRNAs) exhibited typical characteristics of virus in plant hosts. Through comprehensive evolutionary analysis, we demonstrated that SRAV should be a positive single-stranded RNA (ssRNA) flavi-like virus that infects alfalfa, rather than a member of the double-stranded RNA (dsRNA) of the family *Endornaviridae*. Our findings suggest that SRAV represents a unique class of plant-hosted flavi-like viruses with unusual genome organization and evolutionary status, differing from previously identified flavi-like viruses documented to infect plants. The latter shows a close evolutionary relationship to flavi-like viruses primarily found in plant-feeding invertebrates and lacks evidence of triggering host RNA interference (RNAi) responses so far. In summary, our study resolves the taxonomic controversy surrounding SRAV and suggests the potential existence of two distinct clades of plant-hosted flavi-like viruses with independent evolutionary origins. Furthermore, our research provides the first evidence of plant-hosted flavi-like viruses triggering the host’s RNAi antiviral response. The widespread occurrence of SRAV underscores its potential ecological significance in alfalfa, a crop of substantial economic importance.

## Introduction

Continuous advancements in high throughput sequencing (HTS) technologies and bioinformatics methodologies have greatly facilitated the identification of divergent viral sequences that may have been previously overlooked (Charon et al. 2022). As a consequence, some divergent viral contigs encoding “orphan” proteins have been identified and documented (Kuchibhatla et al. 2014). Those divergent viral contigs, presumed to represent divergent viral entities, often pose challenges in their classification within the virus taxonomy lineage, given the lack of sequence similarity homologs. Moreover, several issues warrant careful consideration before interpreting the results from metagenome or metatranscriptome datasets (Togoobat et al. 2023). Do these divergent viral contigs truly represent viruses, or are they instead host genetic sequences harboring endogenous viral elements (EVEs)? Additionally, determining the *bona fide* host of a virus, particularly those viruses identified solely through *in silico* metatranscriptomic or metagenomic analysis, remains an open question.

HTS technologies have not only led to the discovery of numerous new viral sequences but have also provided new insights into virus-host relationships. Taking flavivirids (family *Flaviviridae*) as an example, typical flavivirids within the family *Flaviviridae* are characterized as monopartite, single-stranded, positive-sense RNA viruses between 9 to 13 kilo-bases (kb) in length that code a large polyprotein (Blitvich and Firth, 2015; Parry and Asgari, 2019). Flavivirids are primarily transmitted by arthropods and can cause severe illnesses in vertebrates, including humans, on a global scale (Bamford et al. 2022; Pierson and Diamond, 2020). Most recognized flaviviruses in the family *Flaviviridae* are transmitted horizontally between arthropods and vertebrate hosts, categorizing them as dual-host viruses. However, not all flaviviruses rotate between arthropods and vertebrates; some are specific to vertebrate hosts, while others seem to be restricted to insects (Blitvich and Firth, 2015).

Our knowledge about the host range of flavivirids has been continuously improved due to the widespread usage of HTS technologies. Metatranscriptomic studies have revealed that flavivirids can also be found in a variety of aquatic invertebrates, including sharks, crabs, giant freshwater prawns, and several other species (Dong et al. 2021; Parry and Asgari, 2019). In addition, data mining of publicly available transcriptome datasets revealed that the host range of *pestivirus* (a genus in the family *Flaviviridae*) includes amphibians, reptiles, and ray-finned fish (Mifsud et al. 2022). A research study published in 2022 revealed the presence of an orphan viral genome in alfalfa, which possesses unusual genome organization and was named Snake River Alfalfa virus (SRAV). The authors proposed that SRAV is the first flavi-like virus identified in a plant host (Dahan et al. 2022). However, this viewpoint was later challenged. A study published in 2023 proposed that SRAV should be classified as a member of the family *Endornaviridae* rather than *Flaviviridae* (Postnikova et al. 2023). Moreover, some flavi-like viral sequences encoding a larger polyprotein and possess longer genomes (up to 23 kb in length) compared to classical flavivirids, have also been identified in plants (Atsumi et al. 2013; Schönegger et al., 2022; Debat and Bejerman, 2023). These flavi-like viruses have distinct genome organizations from classical flavivirids but group within the family Flaviviridae in phylogenetic analysis based on the RNA-dependent RNA polymerase (RdRp) domain (Paraskevopoulou et al. 2021). Additionally, these flavi-like viruses with longer genomes identified in plants exhibited a close phylogenetic relationship to those flavi-like viruses mainly found in plant-feeding invertebrates (Dolja, Krupovic and Koonin 2020). However, so far, there is no research support that these flavi-like viruses with longer genomes, documented to infect plants, can activate the host’s RNAi antiviral response. This raises the doubts whether these viruses are truly plant viruses, despite some studies showing that a flavi-like virus (gentian Kobu-sho-associated virus, GKaV) is very widely detected among gentians and other plants (Atsumi et al. 2013; Atsumi et al. 2023; Shaffer et al. 2022).

RNA interference (RNAi) is a conserved antiviral mechanism across eukaryotes (Ding 2010, 2023). In the antiviral RNAi process, virus infection induces Dicer to process virus-specific double-stranded RNA into small interfering RNAs (siRNAs) (Ding 2010, 2023). During this process, the virus-derived small interfering RNAs (vsiRNAs) will be enriched in the hosts, which can be easily detected by deep sequencing (Zheng et al. 2017). Hence, through deep sequencing and bioinformatic analysis of small RNA populations from infected plants, both RNA and DNA viruses, along with viral satellites and viroids, can be identified and their genomes partially or fully reconstructed (Kreuze et al. 2009; Wu et al. 2010; Zheng et al. 2017; Pooggin 2018). Previous research has demonstrated that insect-borne viruses exhibit distinct length distribution patterns in their planthopper vectors compared to rice hosts. This suggests that the vsiRNAs profile can provide insights into the origins of these vsiRNAs (Yang et al. 2018). Moreover, analyzing the distinctive vsiRNA profile can serve as an effective method to differentiate between an endogenous viral element (EVE) and a genuine virus source (Pooggin 2018).

In the present study, we performed a large-scope analysis of 77 publicly accessible small RNA datasets from alfalfa suggesting that SRAV represents a true episomal virus capable of triggering the plant host’s RNAi antiviral defense. Subsequent evolutionary analysis revealed that SRAV is a flavi-like virus infecting alfalfa, rather than a member of the family *Endornaviridae*. In brief, our analysis suggests the potential existence of two distinct types of plant-hosted flavi-like viruses with different evolutionary origins. Additionally, it provides the first RNAi evidence that plants can serve as hosts for flavi-like viruses.

## 2. Methods

### 2.1 Homology search and Conserved RdRp motif identification

The ORF Finder online server (https://www.ncbi.nlm.nih.gov/orffinder/) was used to predict the open reading frame (ORF) of the SRAV viral genome (Accession: ON669064.1). The conserved domains of SRAV polyprotein (Accession: USJ75181.1) were predicted using two methods, NCBI CD-search (https://www.ncbi.nlm.nih.gov/Structure/cdd/wrpsb.cgi) and InterProScan (https://www.ebi.ac.uk/interpro/). In addition, to identify as many functional domains as possible, HHpred was selected to the potential functional domains. HHpred employs a profile-against-profile search method, enabling the detection of distant protein sequence similarities (Söding, Biegert, and Lupas 2005).

To rule out the possibility that SRAV is merely alfalfa gene carrying the endogenous viral element (RdRp), we first compared SRAV with the alfalfa genome (Long et al. 2022) using BlASTn and BLASTx with default parameters.

To investigate whether the SRAV, as an orphan viral genome, harbors the conserved motif in the palm subdomain of RdRp, Geneious software (Kearse et al. 2012) was used to identify and curate the conserved RdRp canonical A, B, and C motifs in the result obtained from multiple sequence alignment. Multiple sequence alignment was carried out using MAFFT, and alignment visualization was achieved using the webserver ESPript 3.0(Robert and Gouet 2014).

### 2.2 Sequence Read Archive datasets collection, processing, and virome analysis

The accession numbers of alfalfa-related small RNA datasets were collected through a comprehensive literature review of research studies involving small RNA sequencing in alfalfa. We first checked the downloaded datasets for small RNA sequencing adapter sequences. If adapter sequences were present, cutadapt (v1.16) (Martin, 2011) was used to remove the adapter sequences. Reads with lengths between 18 nt and 30 nt that contained no ambiguous nucleotides (’N’) were retained as clean reads. The clean reads were then subjected to virome analysis using VirusDetect (Zheng et al. 2017).

### 2.3 SRAV occurrence analysis and vsiRNAs profile characterization

The clean reads were clustered into unique reads using an in-house Python script. Next, the remaining small RNA clean reads were aligned to the SRAV virus reference genome (accession number: ON669064.1) using Bowtie (Langmead et al. 2009), allowing for up to one mismatch. We then analyzed the mapped small RNAs to explore the profile of vsiRNAs, including their length distribution, 5’ terminal nucleotide preference, genome distribution, and polarity.

The average sequencing depth and coverage breadth were calculated as detailed previously (Togoobat et al. 2023). To reduce false positives caused by the random alignment of short fragments, we only considered SRAV in the sample if the vsiRNAs mapped to the SRAV genome met the following criteria: an average sequencing depth of at least 4X and viral genome coverage of more than 60%. Under these conditions, the presence of SRAV was considered confident.

### 2.4 Phylogenetic analysis

Phylogenetic analyses were conducted using multiple sequence alignments (MSA) of either the RdRp domain region or the whole polyprotein sequence across various viral groups. The polyprotein sequences of representative members from different viral groups were retrieved from the NCBI GenBank database. The RdRp domain regions were predicted using a CD-search web server, and the corresponding RdRp domain sequences were extracted based on their positions within the polyprotein or ORF encoding RdRp.

The alignment of RdRp sequences or polyprotein sequences was achieved by Multiple Alignment using Fast Fourier Transform (MAFFT) (Katoh and Standley 2013). The obtained multiple sequence alignments were subjected to trimAl to conduct automated alignment trimming (Capella-Gutiérrez, Silla-Martínez, and Gabaldón 2009). The trimmed MSA files were then subjected to IQ-TREE (v1.6.6) (Nguyen et al. 2015) for phylogenetic tree construction through the maximum likelihood method. The best-fitted model selection was performed by ModelFinder implemented in IQ-TREE (Kalyaanamoorthy et al. 2017), and the confidence in the topology was assessed using 5000 ultra-fast bootstrap replicates.

The sequences of all the RdRP domain regions or polyproteins sequences with accession numbers, coupled with best-fitted model for each tree, can be found in Supplementary Table S2.

## 3. Results

### 3.1 Genome organization and conserved RdRp motif analysis of SRAV

The genome organization of SRAV from two independent research groups are compared and displayed. These two SARV isolates encode polyproteins of the same length (3835 aa), with slight differences in the lengths of the 5’UTR and 3’UTR (Fig. 1A). The BLASTn analysis reveals a similarity of 99.85% (with a query cover of 100%) between these two SRAV isolates, indicating a high degree of sequence agreement. The polyproteins of these two SRAV isolates show a 99.9% similarity in a BLASTp analysis (with 100% query coverage), differing by only four amino acids. The comparison between the SRAV genome and alfalfa genomes using BLASTn reveals no sequence similarity.

**Fig.1.**
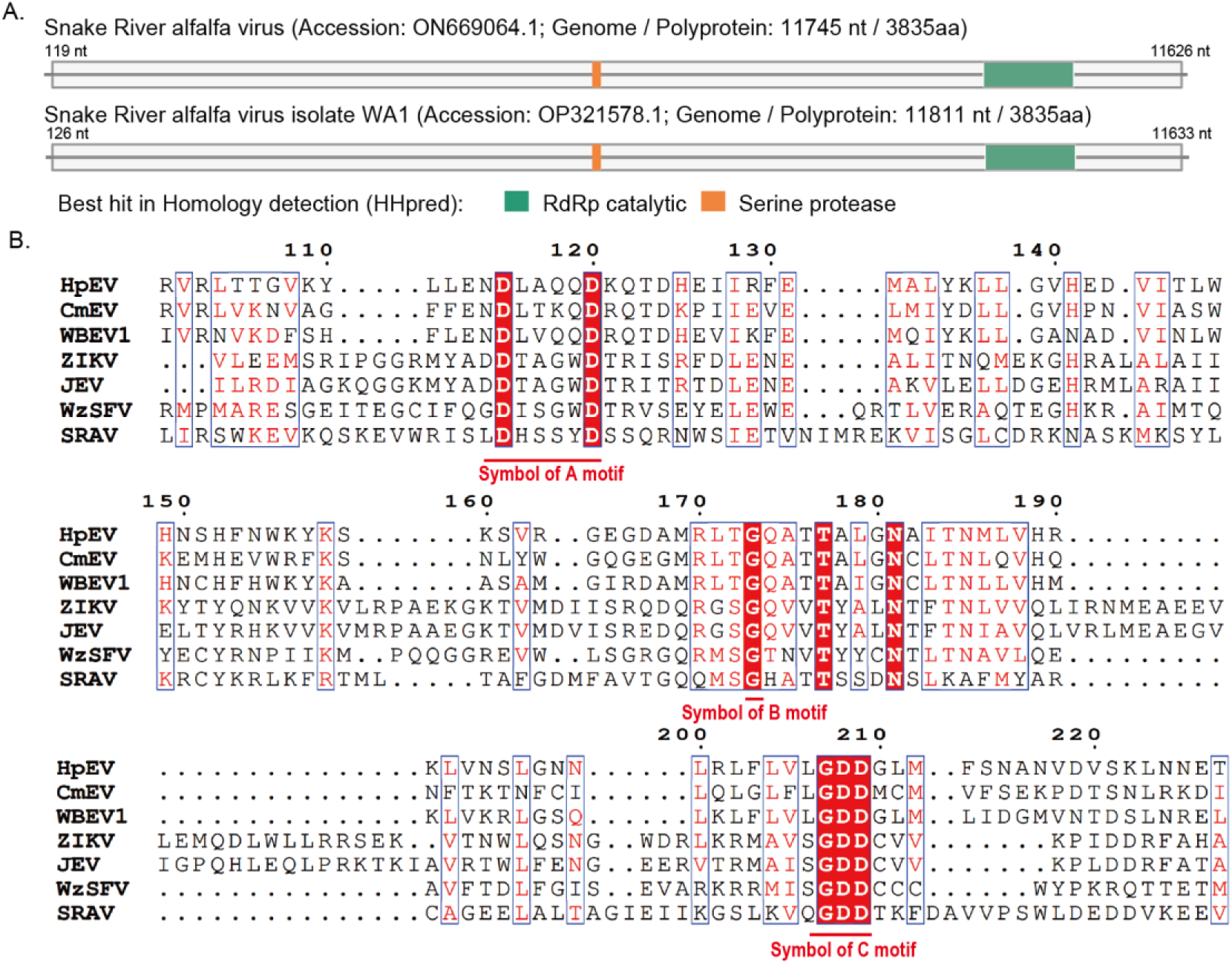
Genome organization of SRAV and multiple alignment of the conserved RdRp palm sub-domain of SRAV and other eukaryotic viruses. (A) Schematic representation of the function domain predicted by HHpred on SRAV. The grey line in the diagram represents the virus genome. Each box represents an open reading frame (ORF) of the virus. Conserved functional domains are color-coded, and the corresponding names are shown on the figures’ right side. (B) Multiple sequence alignment of the conserved RdRp palm domain of SRAV and representative members of the family *Flaviviridae* and *Endornaviridae*. Abbreviations of virus names: HpEV, hot pepper endornavirus; CmEV, Cucumis melo endornavirus; WBEV1 winged bean endornavirus 1; ZIKA, Zika virus; JEV, Japanese encephalitis virus; WzSFV, Wenzhou shark flavi-like virus; SRAV, Snake River alfalfa virus.

Interestingly, upon comparing the SRAV genome sequence with the NCBI NR database for BLASTx analysis, we found no sequence homologs except for SRAV itself. This suggests that SRAV may represent a highly distinct viral sequence encoding an orphan protein.

HHpred analysis indicate that all these two SRAV isolates could be annotated with two functional domains, namely, RdRp catalytic domain and Serine protease domain (Fig. 1A). The annotation detail of function domain annotation, including hit information, E-value or probability, could be found in Fig. S1. It’s worth noting that RdRp catalytic domain can be annotated using all three tools including CD-search, InterProScan, and HHpred. However, the Serine protease domain can only be annotated using HHpred (Fig. S1).

We also analyzed the palm sub-domain of the RdRP protein derived from SRAV and representative members of the *Endornaviridae* and *Flaviviridae* families, which represents the highly conserved catalytic core of RdRP that consists of three motifs: A, B, and C (Fig. 1B). Although SRAV did not find any hits besides itself using NCBI BLASTx search, this orphan virus possesses the typical motifs A, B, and C (Fig. 1B). For instance, similar to representative members of the *Endornaviridae* and *Flaviviridae* families, SRAV exhibits the characteristic Motif A (D-x(4,5)-D), Motif B (a conserved glycine), and the typical C motif (GDD triad) (Fig. 1B).

### 3.2 The widespread occurrence of SARV isolates in alfalfa

To determine whether SRAV is widespread in alfalfa and capable of activating host’s antiviral responses, we conducted a large-scale analysis of all publicly available alfalfa small RNA datasets deposited in the NCBI SRA database. We found that, in each sample, there were varying number of reads that could be mapped to the SRAV genome (ranging from a minimum number of 10 to a maximum of 6370129) (Supplementary Table S2). To avoid false positives caused by the random alignment of small RNA reads, which are typically short, we only consider the presence of SRAV as reliable in samples that pass the criteria we set: an average sequencing depth more than 4X and genome coverage more than 60%). Based on these criteria, we confidently identified the presence of SRAV in 68 out of 77 samples (Fig. 2A). Given that these 77 samples are derived from 10 different research projects (each with distinct BioProject accession numbers), the occurrence of SRAV across these independent studies can be deemed reliable. Furthermore, within four specific BioProjects (SRP110842, SRP336109, SRP064230, and SRP201465), the presence of SRAV was consistently reliable in every sample (Fig. 2A). Based on the meta-information associated with each sample, we found that SRAV can be identified across a diverse range of alfalfa cultivars, indicating its widespread presence among different genetic backgrounds (Fig. 2B). Additionally, SRAV was detected in various tissues of alfalfa, including leaves, stems, roots, seeds, shoots, non-embryogenic, cut cotyledon, and cotyledon embryo, demonstrating its broad tissue distribution within the plant hosts (Fig. 2C). Although SRAV was first identified through metatranscriptome sequencing analysis in 2022, we were able to detect the presence of the virus in small RNA samples dating back to 2014 (Fig. 2D). We also found that SRAV has a wide geographical distribution, being present across the Pacific Ocean in both Asia and North America, primarily in China and the United States (Fig. 2E). It should be noted that this does not indicates that SRAV is absent in other regions. Rather, it indicates that alfalfa from these areas has corresponding high-throughput small RNA sequencing data available for analysis. Additionally, we analyzed the length distribution of SRAV vsiRNAs in the 68 samples in which SRAV could be detected. We found that, without exception, the majority of SRAV vsiRNAs were 21 nt and 22 nt in length, with 21 nt being more abundant (Fig. 2F). This pattern is characteristic of typical plant virus vsiRNAs.

**Fig.2.**
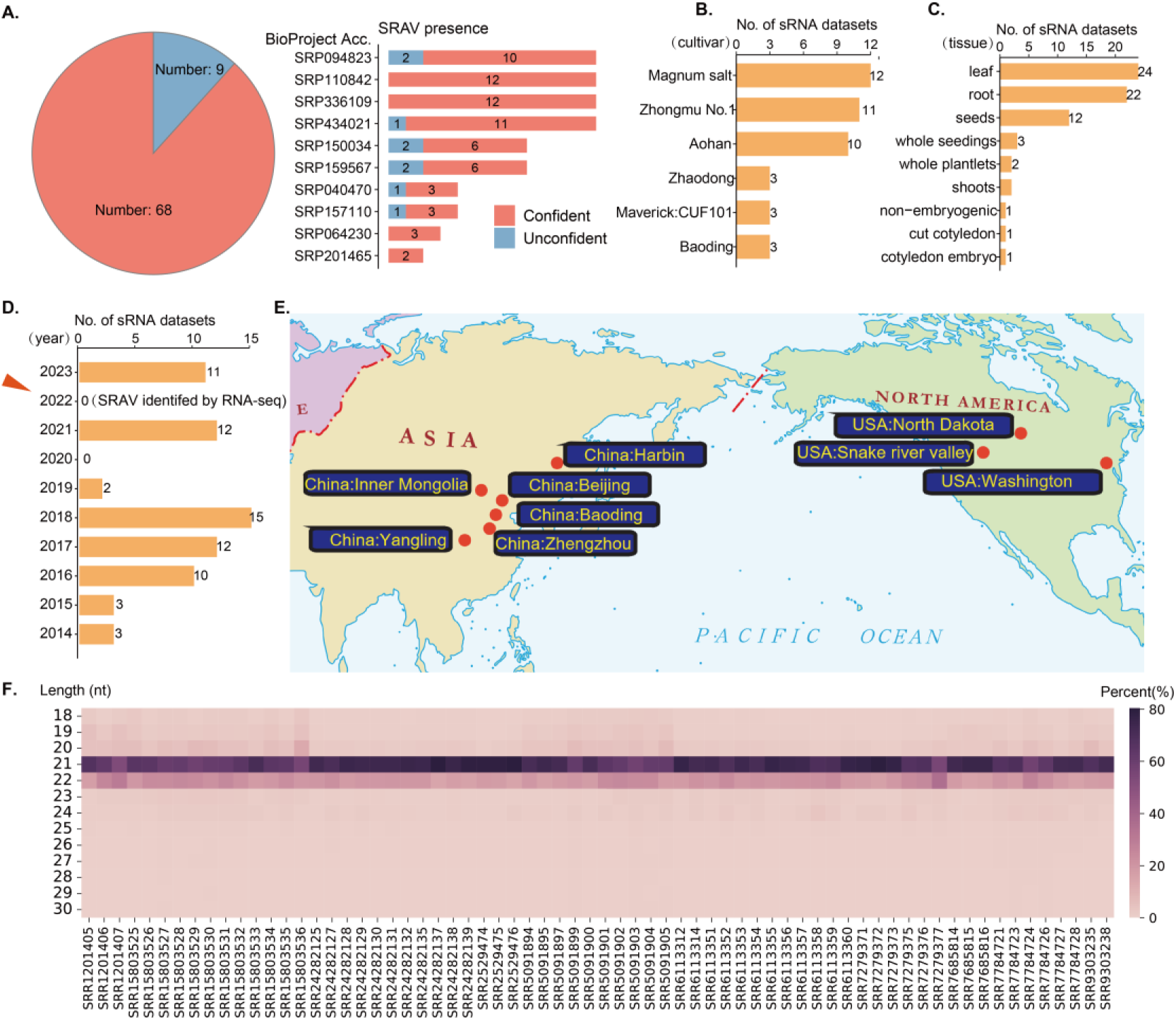
SARV is widespread in Alfalfa and can trigger host’s RNAi antiviral response. (A) The presence of SRAV in each dataset. Confident: the presence of SARV is reliable. Unconfident: the presence of SARV is unreliable. (B) The number of datasets in which SRAV could be reliably detected across different alfalfa cultivars. (C) The number of datasets in which SRAV could be reliably detected across different alfalfa tissue types. (D) The number of datasets in which SRAV could be reliably detected in different years. (E) The currently known geographic distribution of SRAV. (F) The length distribution of SRAV vsiRNAs across different datasets.

### 3.3 Host RNAi antiviral response to SRAV and other RNA viruses

To gain deeper insights into the interaction between SRAV and the host through the RNA interference (RNAi) mechanism, we selected BioProject SRP159567 for a detailed analysis of vsiRNA characteristics for SRAV and other RNA viruses. We chose this BioProject because the proportion of SRAV vsiRNAs is significantly high in some samples within this BioProject (Supplementary Table S2). In the eight samples archived in BioProject SRP159567, the presence of SRAV is considered reliable in six samples. The proportion of SRAV vsiRNAs in the total small RNA population ranges from 0.006% (SRR7784728) to 15.697% (SRR7784727). These six samples, deemed to have a reliable presence of SRAV, encompass various genotypes and tissue types (Fig. 3A). The SRAV vsiRNA profiles of those 6 samples show a high degree of consistency, with 21 nt vsiRNAs accounting for more than 60%, while the proportion of 22 nt vsiRNAs fluctuates around 20% (Fig. 3B). When we analyzed sample SRR7784727 (Shoots sample of the Altet genotype) to study the length distribution, strand preference, and 5’ base preference of vsiRNAs, we found that the SRAV vsiRNAs were relatively evenly derived from both the positive and negative strands of the genome, with a 5’ base preference for nucleotide “U” (uracil) (Fig. 3C). Similarily, we could also observe that the “U” preference of SRAV vsiRNAs is widespread across the majority of alfalfa small RNA samples (Fig. S2A). When analyzing the strand preference of SRAV vsiRNAs, we observed that they can originate from both the sense and antisense strands, typically in similar proportions. However, in samples SRR7784724 and SRR7784726, vsiRNAs from the sense strand exhibited a relatively higher proportion. Notably, both samples were derived from root tissue, suggesting potential variations in the vsiRNA biogenesis process across different tissues in plant hosts (Fig. 3D). Our findings revealed a relatively uniform distribution of vsiRNAs across the entire SRAV viral genome. However, certain localized regions emerged as hotspots for vsiRNA production, with 21 nt and 22 nt vsiRNAs being the predominant types within each region (Fig. 3D-E).

**Fig.3.**
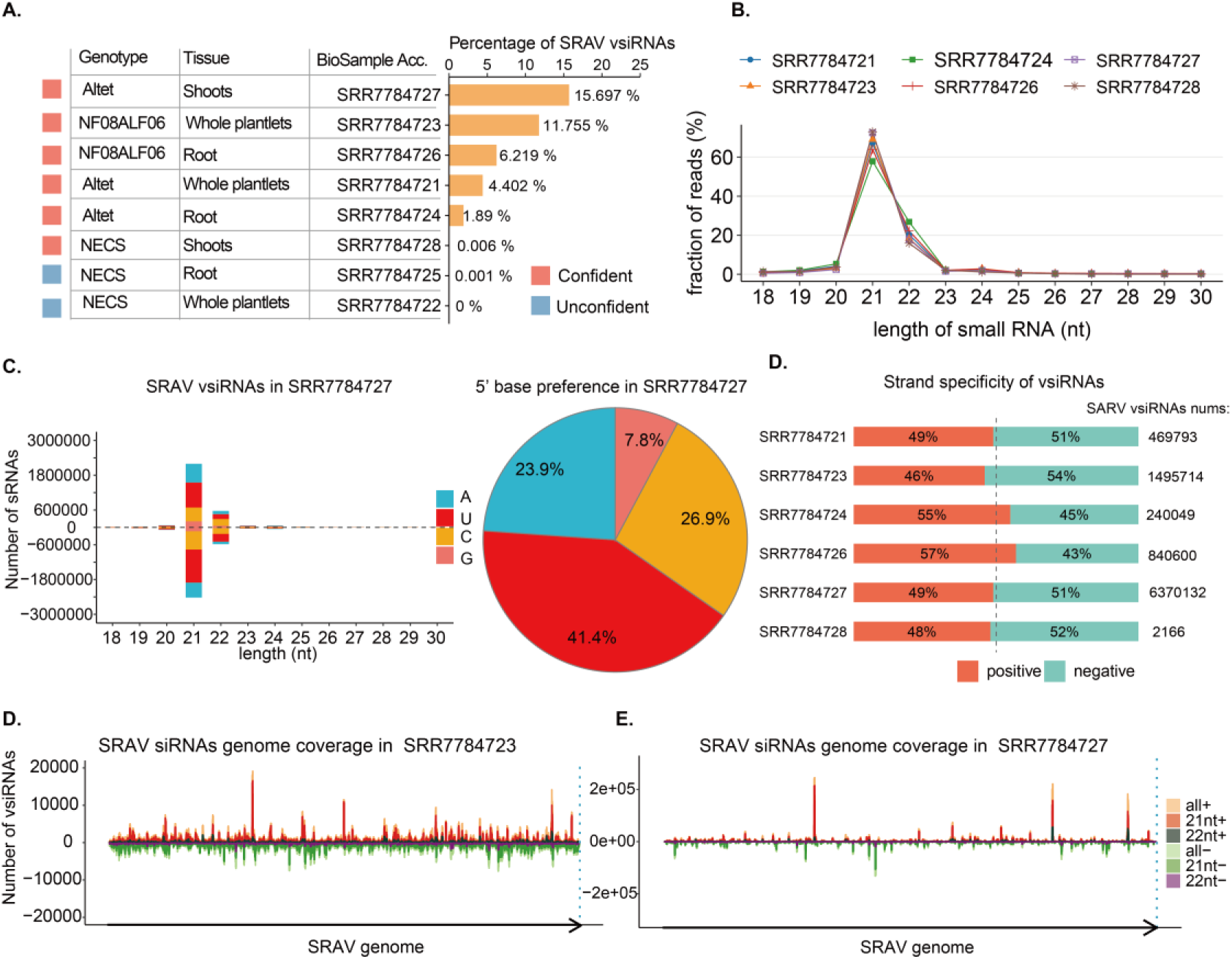
Host RNAi antiviral response to SRAV and other RNA viruses identified in BioProject SRP159567. (A) The metadata information and SRAV vsiRNAs percentage in each samples. (B). length distribution of SRAV vsiRNAs in samples where SRAV was reliably detected. (C). Length distribution and 5’ termini base preference of SRAV vsiRNAs in sample SRR7784727 (D). SRAV vsiRNAs polarity in samples where SRAV was reliably detected. (E) Distribution of SRAV vsiRNAs alongside the viral genome in SRR7784723. (F) Distribution of SRAV vsiRNAs alongside the viral genome in SRR7784727.

We conducted an in-depth viromic analysis of small RNA datasets obtained from BioProject SRP159567 using VirusDetect. Apart from identifying SRAV, as shown in Fig. S2B, we also detected three other alfalfa viruses: Medicago sativa amalgavirus 1, Medicago sativa alphapartitivirus 1, and Medicago sativa alphapartitivirus 2, as detailed in Fig. S2C. The length distribution patterns of SRAV vsiRNAs closely resembled those of the other three alfalfa viruses (Fig. S2D), suggesting that SRAV is indeed a virus infecting alfalfa, capable of triggering the plant’s antiviral RNAi response.

### 3.4 *Flaviviridae* or *Endornaviridae*? The phylogenetic status of SRAV

After confirming that SRAV is a plant virus, we investigated its evolutionary status. There is ongoing academic controversy regarding whether SRAV belongs to the family *Flaviviridae* or *Endornaviridae* (Dahan et al. 2022; Postnikova et al. 2023). Polyprotein sequences of several representative members of the family *Flaviviridae* and *Endornaviridae* were retrieved and aligned. After the alignment sequence trimming, a maximum likelihood tree with best-fitted substitution model was constructed. The resulting phylogenetic analysis revealed that SRAV clusters within the family *Flaviviridae* with a strong bootstrap support (Fig. 4). However, this taxonomic classification contradicts the conclusions of most recent research about SRAV evolutionary status (Postnikova et al. 2023), which classified SRAV as a member of the *Endornaviridae* family. To address the potential influence of differing sequence sets used in phylogenetic tree construction, we reconstructed the evolutionary tree using the same sequence sets from the previous study (Postnikova et al. 2023). Despite this, our results still clearly indicate that SRAV clusters within the family *Flaviviridae*, as shown in Fig. S4.

**Fig.4.**
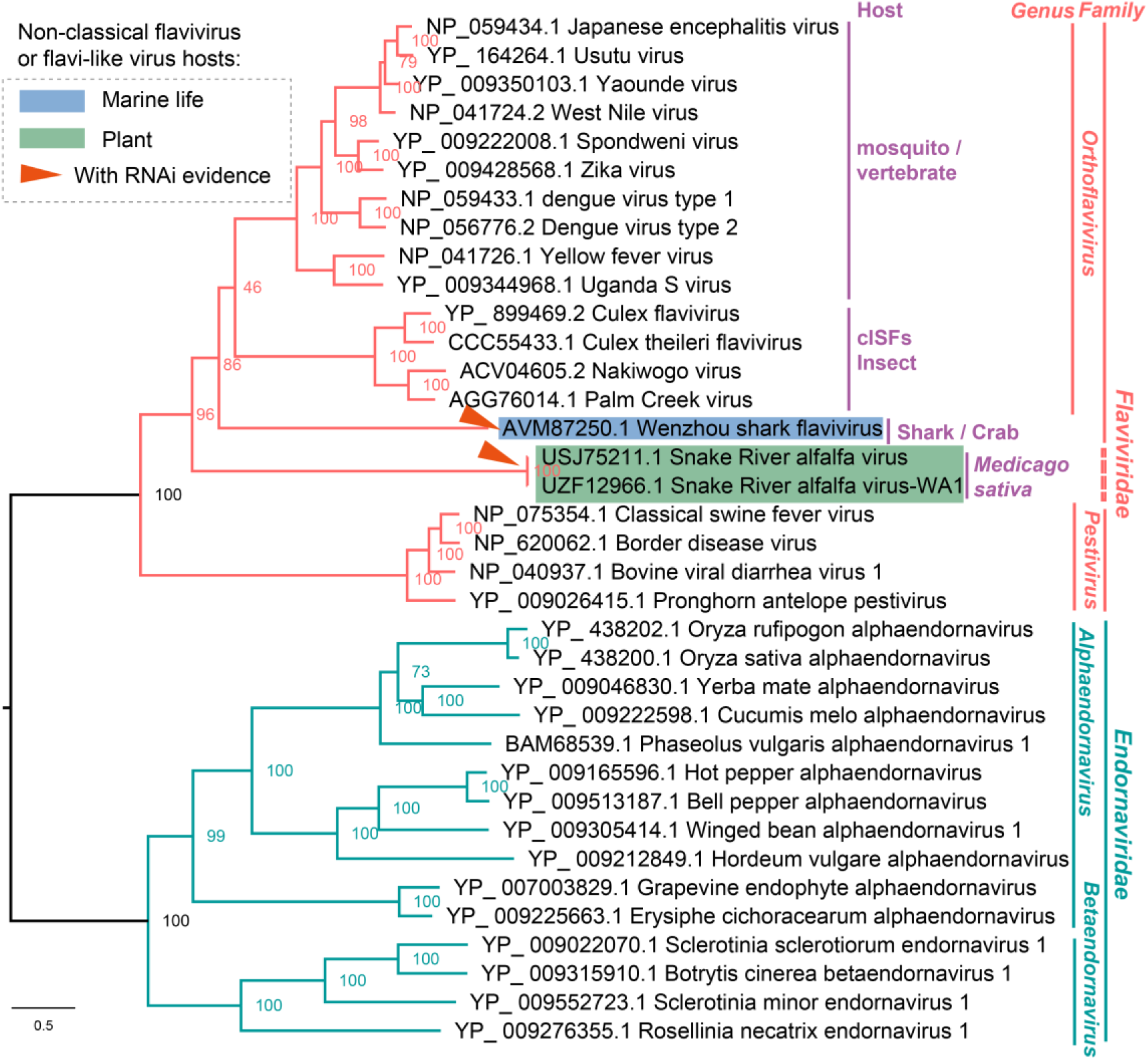
Maximum likelihood (ML) phylogenetic tree from the polyprotein of SRAV and representative members of family *Flaviviridae* and *Endornaviridae*. The clades in red refer to the *Flaviviridae*, and the clades in green refer to family *Endornaviridae*. cISF: classic insect specific flaviviruses. The taxa represented by the red triangle are viruses that can activate the host’s RNAi antiviral response. Sequences used for this tree are listed in Supplementary Table S1.

Based on the evolutionary analysis so fa, it is premature to definitively conclude that SRAV belongs to the family *Flaviviridae*. Given that SRAV represents an orphan viral genome without sequence similarity homologs, the possibility that it belongs to neither family *Flaviviridae* nor *Endornaviridae* cannot be ruled out. Phylogenetic methods might group SRAV and family *Flaviviridae* together because of artificial similarity driven by long branch attraction (LBA) (Susko and Roger 2021). As members of the family *Endornaviridae* are dsRNA viruses, while SRAV is presumed to be a positive ssRNA virus, selecting another group of ssRNA viruses (such as the family *Potyviridae*) instead of the family *Flaviviridae* would still likely result in SRAV clustering with that viral group with well-supported bootstrap value due to LBA (Fig. S4).

To provide more reliable evidence for the evolutionary classification of SRAV, we expanded our analysis by adding a diverse set of taxa. The RdRp domain sequences of representative members from various positive-stranded RNA virus super-families, negative-stranded RNA viruses, as well as members of the SRAV and *Endornaviridae* families, were used to construct the phylogenetic tree. The results indicate that SRAV clusters within the clade of the family *Flaviviridae* group (ssRNA+ flavi-like), supporting its association with this family (Fig. 5). Additionally, we constructed an evolutionary tree by combining the RdRp domain sequence of SRAV with those of representative members from all known polyprotein-encoding viral families. The results consistently showed that SRAV clustered with members of the *Flaviviridae* family (Fig. 6). We further constructed an evolutionary tree using the RdRp domain sequences from SRAV and representative members of different orders within the phylum *Kitrinoviricota*. The resulting phylogenetic tree indicated that SRAV could also be clustered within the order *Amarillovirales*. Since the family *Flaviviridae* is the only family within this order, these findings further support the close evolutionary relationship between SRAV and the family *Flaviviridae* (Fig. S5).

**Fig.5.**
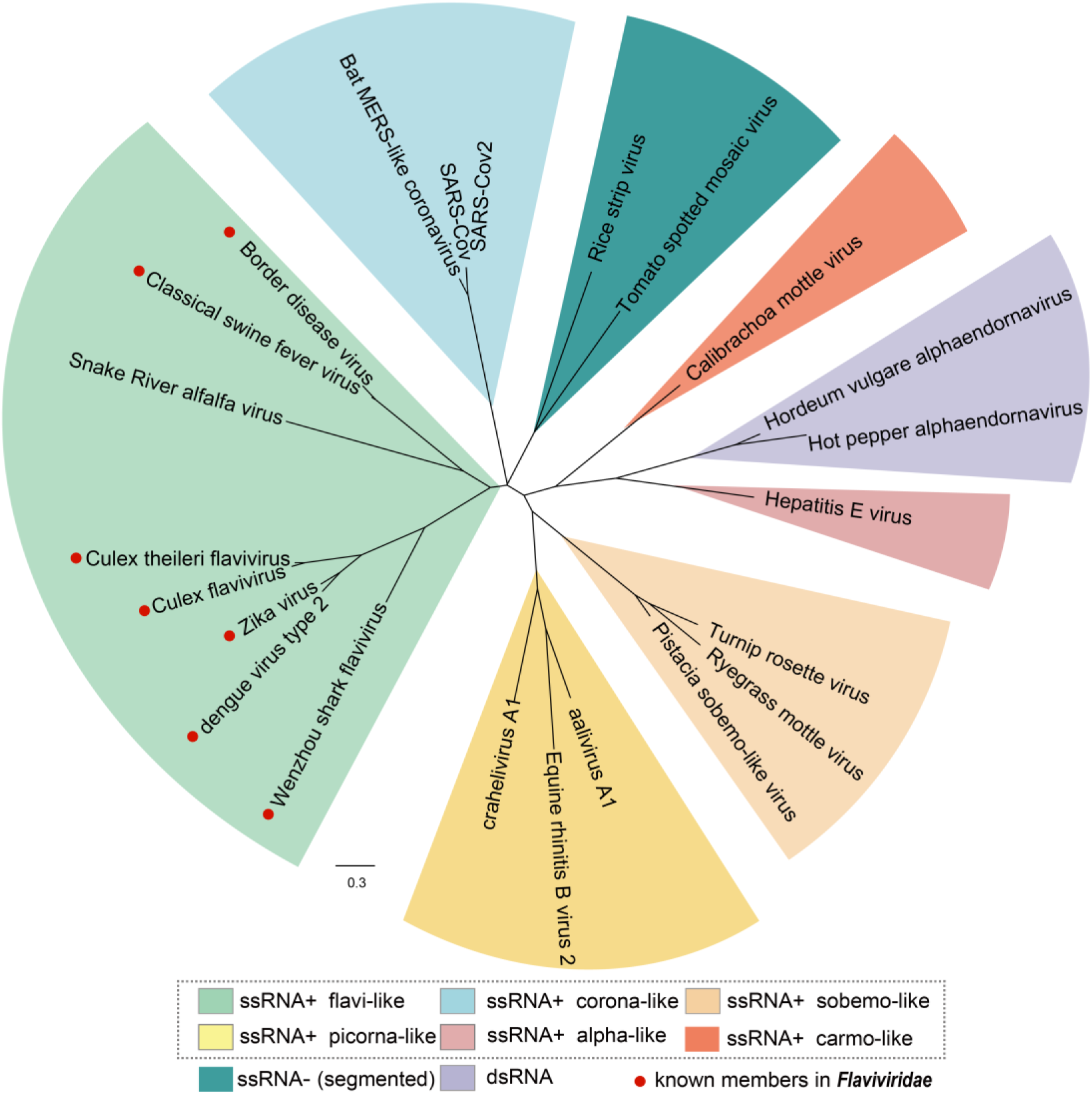
Maximum likelihood (ML) Phylogenetic tree from the RdRp domain of SRAV and representative members of different RNA superfamily. The red points represent the known members in family *Flaviviridae*.

### 3.5 SRAV is evolutionary distant to the flavi-like viruses documented to infect plants

Previous research has identified flavi-like viruses with atypical genome organization in plants, featuring genome lengths twice as long as those of classical flaviviruses. To explore this further, we first compared the polyprotein lengths of different flavivirids groups. We found that the previously identified plant flavi-like viruses have similar polyprotein lengths to flavi-like viruses found in plant-feeding invertebrates. However, the polyprotein of SRAV exhibited a length similar to that of classical flavivirids. This huge difference in polyprotein length suggests that SRAV may have a significant evolutionary distance from the plant viruses documented to infect plants (Fig. 7).

To analyze the evolutionary differences between SRAV and currently published plant flavi-like viruses, a maximum likelihood phylogenetic tree was constructed based on the multiple sequence alignment of the RdRp domain region across different flavivirids groups and the representative members of the family *Endornaviridae*. Notably, this evolutionary tree also incorporates flavivirids identified in marine organisms from a recent research. The phylogenetic tree reveals that flavi-like viruses documented to infect plants thus far are most closely related to flavi-like viruses primarily identified in plant-feeding invertebrates, forming a monophyletic clade. While confirming SRAV’s evolutionary placement within the family *Flaviviridae* once again, the evolutionary tree also underscores that SRAV and the known plant-infecting flavi-like viruses clearly belong to distinct evolutionary clades. Furthermore, the evolutionary statuses of flavi-like viruses identified in marine life are consistent with the previous research (Parry and Asgari, 2019) (Fig. 8).

## 4. Discussion

### 4.1 Orphan viral contigs hidden in metagenomic or metatranscriptomic datasets

Land plants harbor a rich and diverse virome, predominantly composed of RNA viruses, while also featuring significant contributions from reverse-transcribing and single-stranded (ss) DNA viruses (Dolja, Krupovic and Koonin 2020). Consequently, metatranscriptome sequencing emerges as a powerful approach to study the composition and diversity of virome in land plants. However, the current identification of RNA viruses primarily depends on sequence similarity-based searches, which make completely new viruses with extremely low similarity remain undetectable (Charon et al. 2022).

However, some studies have demonstrated that orphan viral contigs can be discovered by bypassing sequence similarity searches and instead leveraging the inherent properties of the viruses. Using the characteristic vsiRNAs profile and high abundance feature, the researchers successfully identified a missing orphan viral segment from a multiple-segment genome (Lu et al. 2022). A clade of divergent viruses represented by orphan viral contigs were identified in oomycetes metatranscriptomic datasets by leveraging the feature of virus dsRNA accumulation (Forgia et al. 2024). Furthermore, conservation within the untranslated regions (UTRs) of certain viruses possessing segmented or multipartite genomes was employed to search for potential orphan viral segments (Zhang et al. 2024).

Thanks to the continuous advancements in HTS technology and bioinformatics analysis methods, some orphan divergent genomes encoding orphan proteins without sequence similarity homologs have been identified (Kuchibhatla et al. 2014). As demonstrated by SRAV (Fig. 1), these sequences may not find any homolog through sequence-to-sequence-based methods like BLAST (Altschul et al. 1990). However, by using methods based on sequence-to-profile or profile-to-profile search, some distant functional domains of those orphan viral genomes can be identified (Kuchibhatla et al. 2014). But the question then arises, that is, how to determine the evolutionary status of these highly divergent viral sequences lacking homologs and how to confirm that these divergent viral contigs represent real viruses rather than endogenous viral elements which is widespread in eukaryotic genomes. In general, comparing candidate viral contigs suspected to be host EVEs with the host genome sequence can often provide some clues. However, this approach necessitates a fully assembled host genome for reference.

4.2 SRAV represents a *bona fide* virus that could trigger the host’s antiviral response in alfalfa Unlike some viral genomes that encode multiple ORFs, such as those from families like *Flexiviridae* which encodes characteristic triple gene blocks (Weng et al. 2024), orphan viral genome with only one ORF encoding a polyprotein have a genetic organization much similar to host genes. In this case, even if the orphan viral genome contains annotated virus-related domains, it is insufficient to determine that it is an episomal virus rather than a host gene containing EVE fragments based on this information alone.

Additionally, a previous study found that some new clades of orphan viral genomes lack the classical catalytic triad motifs in the RdRp protein (Forgia et al. 2022). To determine whether SRAV’s RdRp contains the complete RdRp motif and any non-classical catalytic triads, we conducted a conserved motif analysis on SRAV’s RdRp domain region. We found that, despite SRAV’s RdRp protein not showing detectable similarity to existing ones, it contains conserved motifs A (D_X2-4_D), motif B (conserved G), and motif C (GDD), similar to other eukaryotic viruses. This supports the idea that SRAV represents a true virus (Fig. 1).

However, the best hits for two known viral domains of SRAV are the “Serine protease” of Japanese encephalitis virus (with a probability of 77.9% and an E-value of 6.3e^-5^) and the “RdRp catalytic core” of Classical swine fever virus (with a probability of 98.7% and an E-value of 1.3e^-12^), respectively. Both of these viruses are flavivirids belonging to the family *Flaviviridae*, which primarily infect invertebrates and vertebrates such as mosquitoes, zoonotic animals, and birds. Insects may introduce insect-specific viruses (ISVs) into plant tissues through feeding activities, which could lead to misleading conclusions when these ISV genomes are detected in plant transcriptomic datasets. Furthermore, SARV could also be identified from plant-feeding thrips *Frankliniella occidentalis* (Dahan et al. 2022; Postnikova et al. 2023). Therefore, whether SRAV represent a real plant virus has become a new question.

Evolution has equipped plants with defense mechanisms to combat viral infections (Boualem, Dogimont and Bendahmane 2016). In brief, upon infection, RNA silencing is initiated by the recognition of viral dsRNAs or partially double-stranded hairpin RNAs. These viral RNAs are processed into vsiRNAs through the host’s small RNA biogenesis pathways (Boualem, Dogimont and Bendahmane 2016; Yang et al. 2018). Then we conducted a large scope alfalfa small RNAs datasets analysis to find evidence that SRAV could trigger plant host’s antiviral response. We found that SRAV is detectable in various tissues and cultivars of alfalfa and exhibits a broad geographical distribution (Fig. 2B-E). The profile of SRAV-derived small interfering RNAs (vsiRNAs), including their length distribution and 5’ termini preference, displayed the typical characteristics of plant virus vsiRNAs (Fig. 2, Fig. 3, Fig. S2). It is crucial to highlight that the length distribution of SRAV vsiRNAs is primarily at 21 nt and 22 nt, with 21 nt being more abundant. This characteristic length distribution is conserved among plant virus vsiRNAs and is supported by the presence of similar patterns in other alfalfa viruses (such as Medicago sativa amalgavirus 1) identified in this study (Fig. 2, Fig. 3, Fig. S2).

### 4.3 The evolutionary status of SRAV

Having established that SRAV represents a real plant virus capable of eliciting host antiviral responses, we then investigate its evolutionary status. Determining the evolutionary status of orphan viral genomes remains challenging due to the absence of detectable sequence similarity to any known sequences. Using SRAV as a case study, it was first identified in alfalfa in 2022 and the researchers proposed it as the first plant flavi-like virus within the family *Flaviviridae* (Dahan et al., 2022). However, a subsequent study in 2023 by another research group suggested reclassifying SRAV under the family *Endornaviridae* instead *Flaviviridae* (Postnikova et al., 2023).

The main arguments of the article questioning SRAV’s evolutionary placement are based on several key points. Firstly, SRAV lacks the helicase domain typically found in the family *Flaviviridae* but does contain a poly(A) tail, which is not characteristic of family *Flaviviridae*. Additionally, SRAV is widely distributed in alfalfa, a biological characteristic more similar to viruses in the *Endornaviridae* family. The authors also constructed an evolutionary tree using Maximum Likelihood method using MEGA, which also showed that SRAV is “closer” to the *Endornaviridae* family. Furthermore, the authors conducted agarose gel electrophoresis and observed the presence of dsRNA of the approximately same size of SARV (Postnikova et al., 2023). However, the arguments supporting these challenges are not sufficiently convincing. Since only one RdRp domain could be detected using CD-search or InterProScan in SRAV, an orphan viral genome without any detectable similarity to known sequences, it is highly possible that SRAV contains an unknown helicase domain rather than lacking a helicase domain entirely. As for containing a poly(A) tail, it could be explained by the polymorphism and diversity of the virus, especially for viruses identified across different host species. The observation of dsRNA bands in agarose gel electrophoresis is unsurprising, as single-stranded RNA viruses can also be detected as replicative intermediates (Roossinck, Martin and Roumagnac 2015). The most significant issue is that they made a critical error in their presentation and explanation of the evolutionary tree.

We first constructed a maximum likelihood tree with best-fitted substitution model based on the polyprotein sequences of several representative members of the family *Flaviviridae* and *Endornaviridae*. The phylogenetic tree clearly indicated that SRAV clusters within the family *Flaviviridae* (Fig. 4). Even when utilizing the same dataset as the article challenging SRAV’s taxonomic status to construct an evolutionary tree, it consistently indicates that SRAV belongs to the family *Flaviviridae* (Fig. S3).

To avoid potential misleading interpretations resulting from the long-branch attraction effect during the phylogenetic tree construction, we incorporated a diverse of virus taxa to disrupt those potential long branches. We first added representative RNA virus from different RNA virus supergroup proposed in a literature (Buck 1996), the phylogenetic tree clearly indicates that SARV clusters in “ssRNA+ flavi-like” supergroup instead of dsRNA clade represented by two endornaviruses (Fig. 5).

We expanded our analysis by adding different taxa based on virus genome organization characteristics. We selected representative members from families known to encode polyproteins according to International Committee on Taxonomy of Viruses (ICTV) classification. Subsequently, we constructed an evolutionary tree using the RdRp domain of these viruses alongside the RdRp domain of SRAV. The phylogenetic analysis result also revealed that SRAV and family *Flaviviridae* formed a monophyletic clade (Fig. 6). Additionally, we constructed another evolutionary tree using the RdRp domain sequences from SRAV and representative members of various orders within the phylum *Kitrinoviricota*. The phylogenetic analysis also supported the conclusion that SRAV clusters within the family Flaviviridae, which is the sole family in the order *Amarillovirales* (Fig. S5).

**Fig.6.**
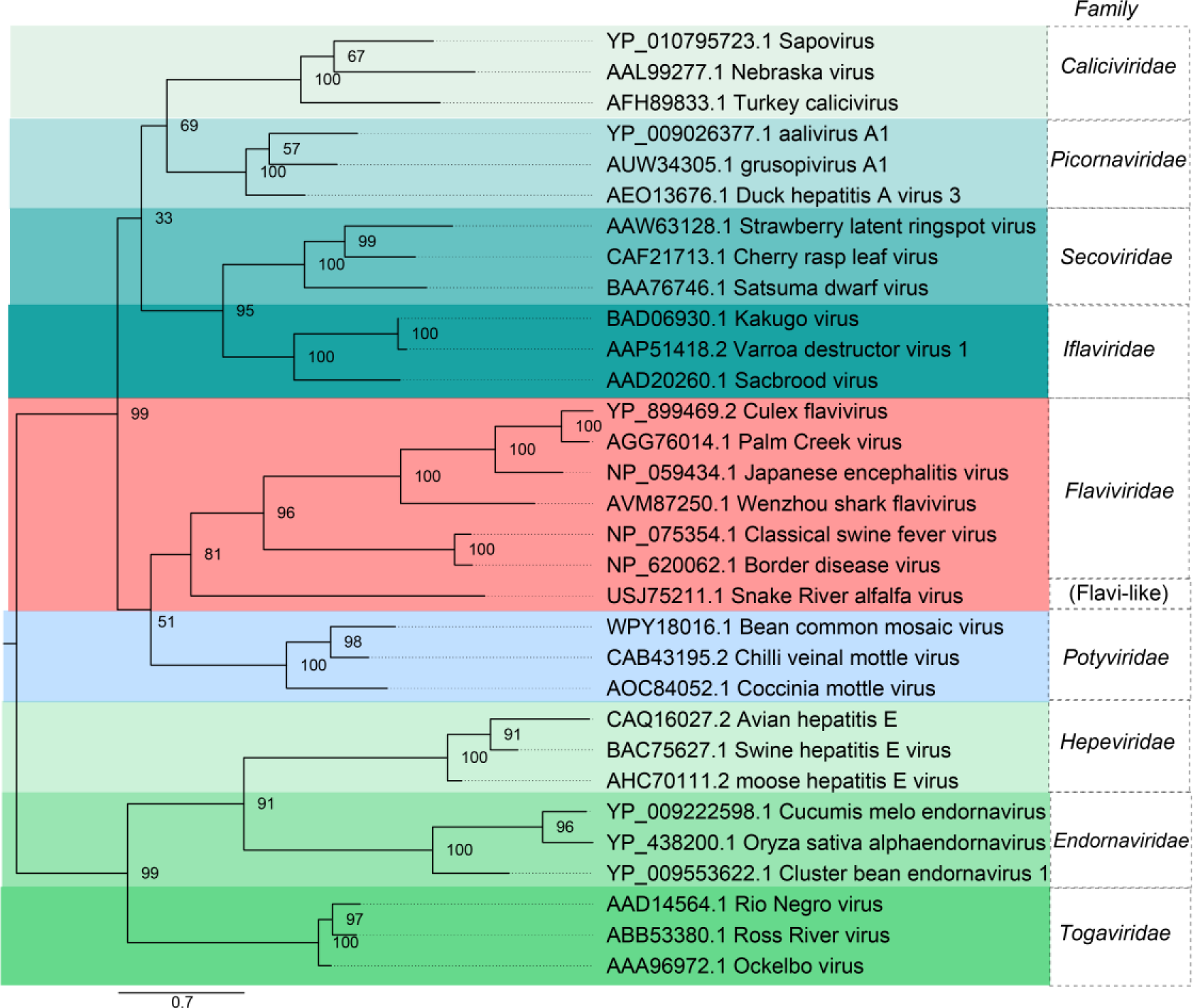
Maximum likelihood (ML) Phylogenetic tree from the polyprotein of SRAV and representative members of different viral family encoding polyprotein.

At this point, the debate regarding the evolutionary status of SRAV becomes clear. It is apparent that SRAV exhibits a closer evolutionary relationship to the family *Flaviviridae* (ssRNA) than to the family *Endornaviridae* (dsRNA). Since the length of the bar below the phylogenetic tree represents the “site substitution rate”, the branch length in the tree can be used to infer the evolutionary distance between different taxa, as proposed by Yang (Yang 1998). From the phylogenetic tree, the branch length distance of the sub-clade composed by SRAV isolates and other sub-clade is considerably long, indicating the substantial evolutionary distance between SRAV with other members of the family *Flaviviridae*. Based on our phylogenetic analysis, it is reasonable to classify SRAV as a distant member of the family *Flaviviridae* and name it a flavi-like virus. As SRAV genome is too divergent to detect similarity to any known sequence, it is conceivable to suggest that SRAV may ultimately serve as the funder of a new genus or even a higher taxonomic category within the broader hierarchy of flavi-like viruses.

### 4.4 SRAV represents a different lineage of plant-hosted flavi-like viruses

In the research work in which SRAV was firstly identified in 2022, the author proposed the SRAV to be the first plant-flavi like viruses (Dahan et al. 2022). However, as early as 2013, a research work reported the identification of a plant flavi-like virus (gentian Kobu-sho-associated virus, GKaV) in gentian with unusual genome organization (Atsumi et al. 2013). Another novel plant flavi-like virus, carrot flavi-like virus 1, which shares its closest genetic similarity with GKaV, was discovered in populations of wild carrots (Schönegger et al., 2022). Subsequently, a research work introduced two novel plant flavi-like viruses by mining of publicly NCBI SRA datasets from sonchus and coptis samples (Debat and Bejerman 2023).

However, the genomes length of the four documented plant flavi-like viruses with unusual genome organization are generally nearly twice as long as those of classic flaviviruses, and their genome length is similar to that of flavi-like viruses identified primarily in plant-feeding invertebrates (Fig. 7). Furthermore, the phylogenetic tree illustrates that these four flavi-like viruses documented to infect plants are closely related to those flavi-like primarily identified in plant-feeding invertebrates, forming a monophyletic clade (Fig. 8). Hence, the possibility that these four flavi-like viruses documented to infect plants are essentially flavi-like viruses of plant-feeding invertebrates introduced into plants through feeding or other behaviors, and subsequently detected accidentally in plant transcriptomic datasets, cannot be neglected. Additional evidence, such as infectious clones, virus inoculation experiments, or the induction of the host’s antiviral response, could provide further insights into whether these four viruses are indeed plant viruses. However, such evidence is currently unavailable for these four plant flavi-like viruses.

**Fig.7.**
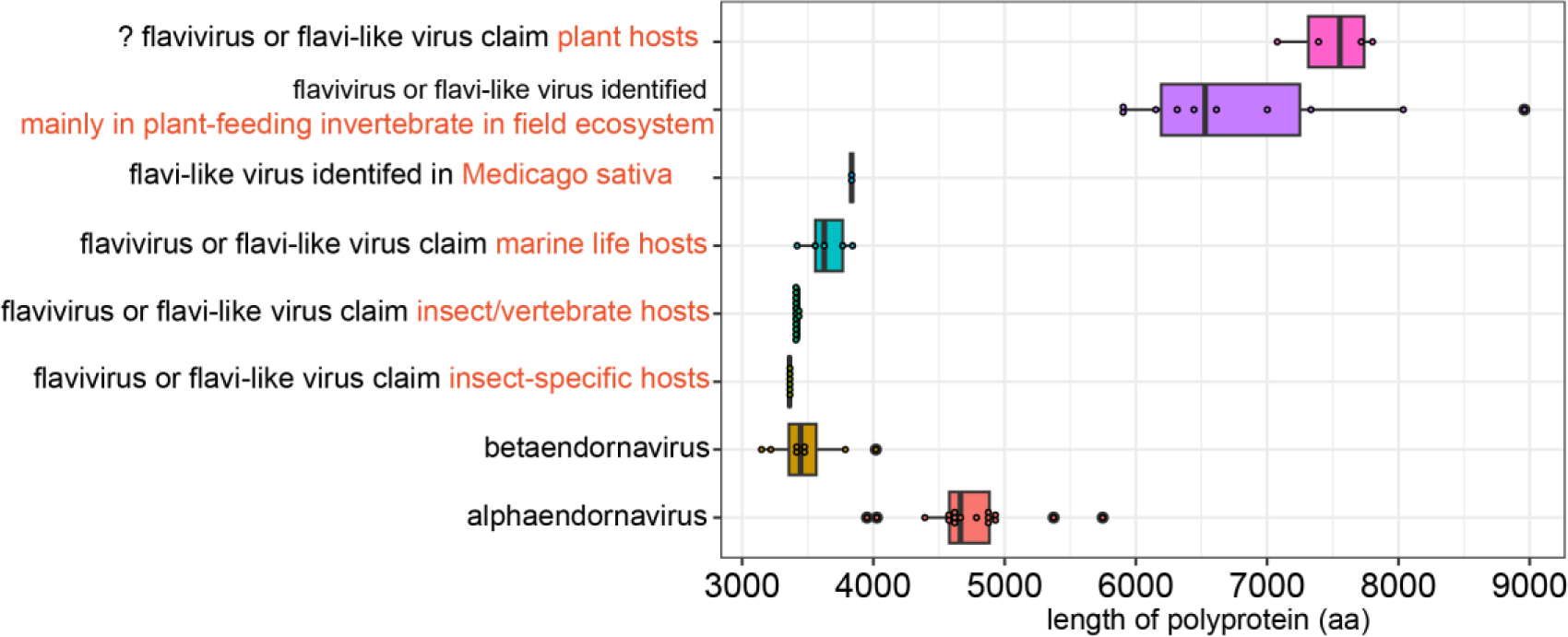
Comparison of polyprotein length of different viral groups.

**Fig.8.**
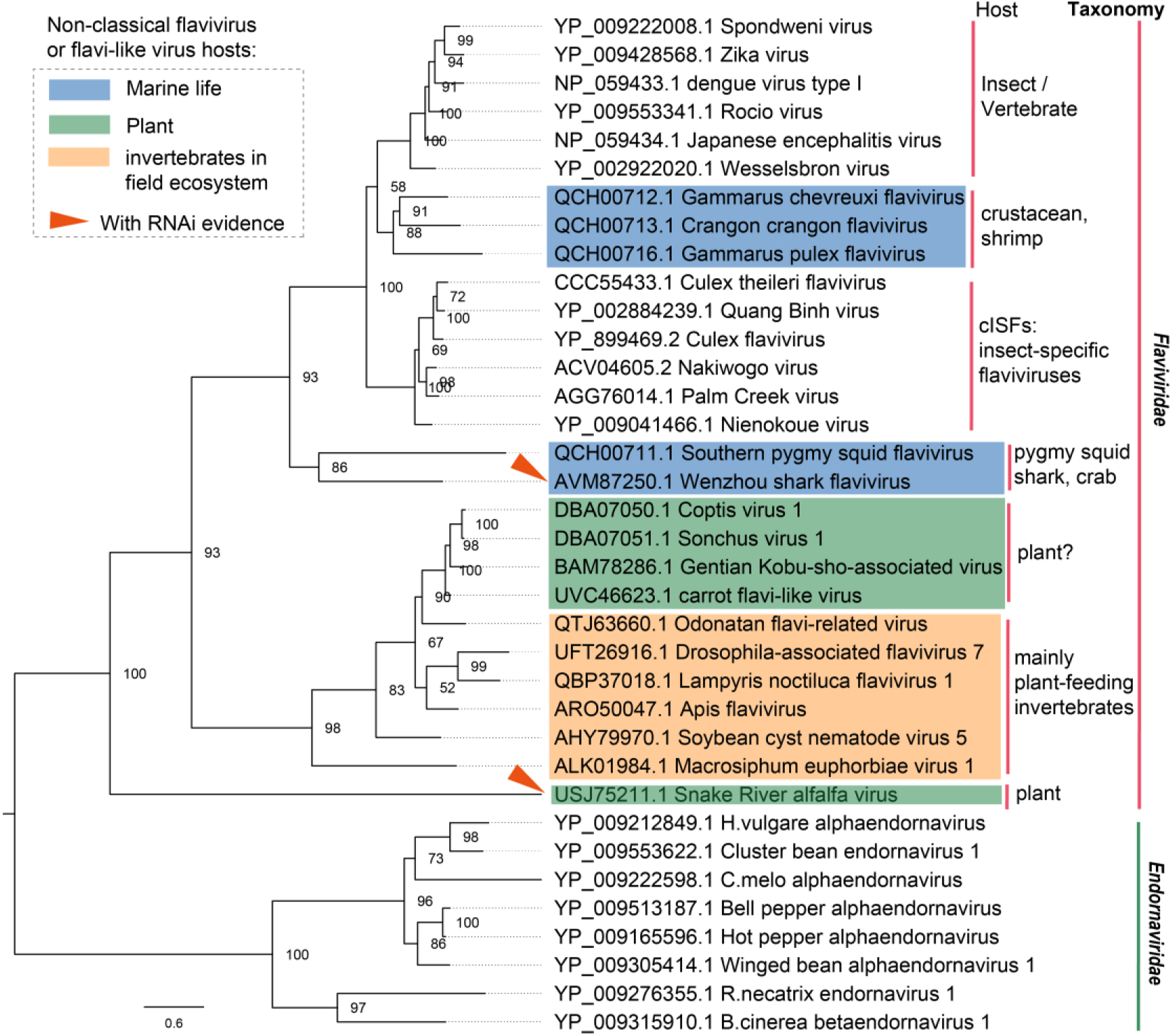
Maximum likelihood (ML) Phylogenetic tree from the RdRp domain of SRAV representative members of family *Flaviviridae* and *Endornaviridae*, including those flavi-like viruses documented to infect plant (green area). The flavi-like viruses with long genome identified in plant-feeding invertebrate were highlighted in yellow area. The flavi-like viruses identified in marine life were highlighted in blue area. The taxa represented by the red triangle are viruses that can activate the host’s RNAi antiviral response.

For comparison, SRAV exhibits a genome length similar to that of classical flavivirids (Fig. 7). While the SRAV genome diverges significantly, making it challenging to detect many known domains, the two domains identified by HHpred exhibit a genome organization similar to that of classic flaviviruses (Dahan et al. 2022). The phylogenetic analysis indicates that SRAV is evolutionary distant to the clade of plant flavi-like viruses with large genome. Although we have questioned whether the four documented plant flavi-like viruses represent real plant viruses, we cannot deny that they are not plant viruses based solely on sequences. As a review paper have discussed, it seems possible that this group of plant flavi-like virus were relatively recently transferred from invertebrates to plants and are undergoing active adaptation to the new host (Dolja, Krupovic and Koonin 2020).

Based on the results of functional domain annotation, genome organization, and phylogenetic analysis, we propose that SRAV represents a distinct lineage of plant-hosted flavi-like viruses. Furthermore, we provide evidence that plant flavi-like viruses could trigger the plant host’s antiviral response for the first time.

## Data availability

Pipeline and scripts used in small RNA datasets processing and multiple sequence alignment files used to construct the phylogenetic tree can be accessed on the GitHub page: https://github.com/BiocompZTXu/project_plant_flavi-like_virus. The publicly available small RNA datasets analyzed in this study are from the BioProject as listed: SRP040470, SRP064230, SRP094823, SRP110842, SRP150034, SRP157110, SRP159567, SRP201465, SRP336109, and SRP434021.

## Acknowledgments

The authors acknowledge the researchers who have generously shared their sequencing datasets with the academic community. This work was supported by National Natural Science Foundation of China (U20A2036). The author would like to express gratitude for the insightful discussions on data analysis provided by Mang Shi from Sun Yat-sen University, as well as Zhen He from Yangzhou University.

---

**Fig. S1.**
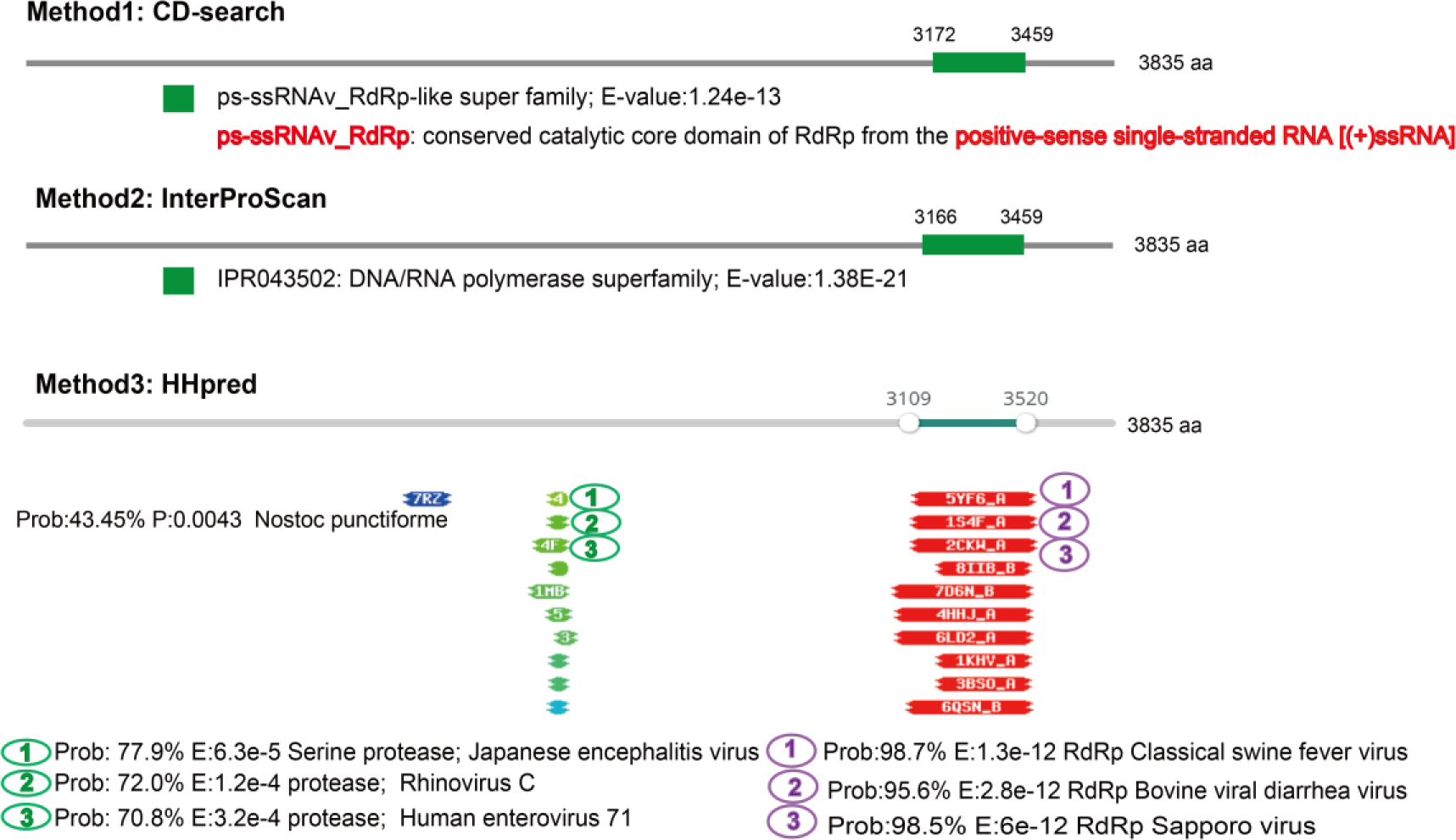
Functional domain predicted by CD-search, InterProScan, and HHpred. The grey line in the diagram represents the virus genome. Box represents domain region.

**Fig. S2.**
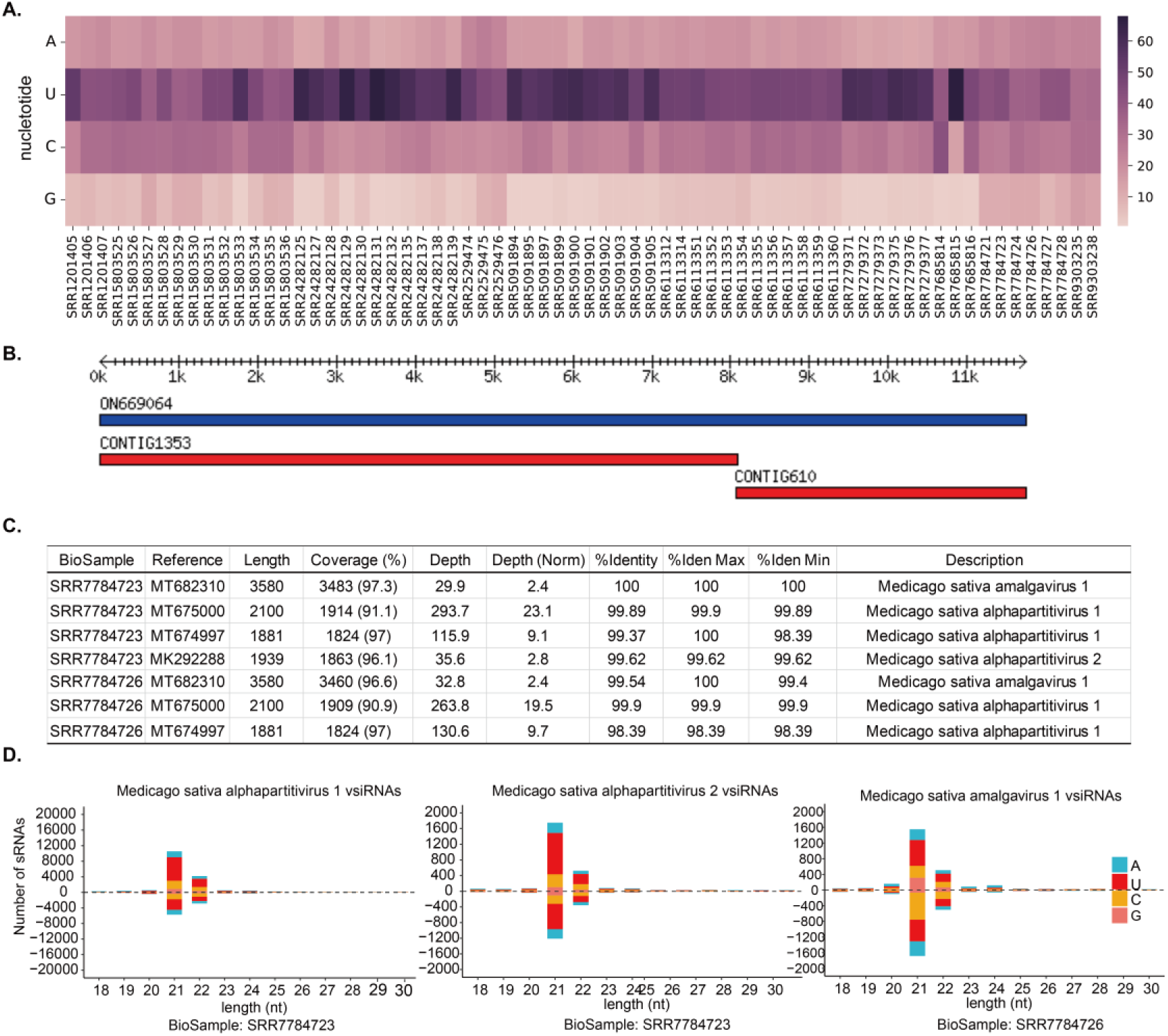
SRAV vsiRNAs characterization and alfalfa small RNA datasets viromic analysis. (A) The 5’ terminal preference of SRAV vsiRNAs across different datasets. (B) Display of assembled contigs aligned to SRAV genome. (C) Viromic analysis results of datasets in BioProject SRP159567. (D) Length distribution and 5’ terminal nucleotide preference of other three alfalfa viruses.

**Fig. S3.**
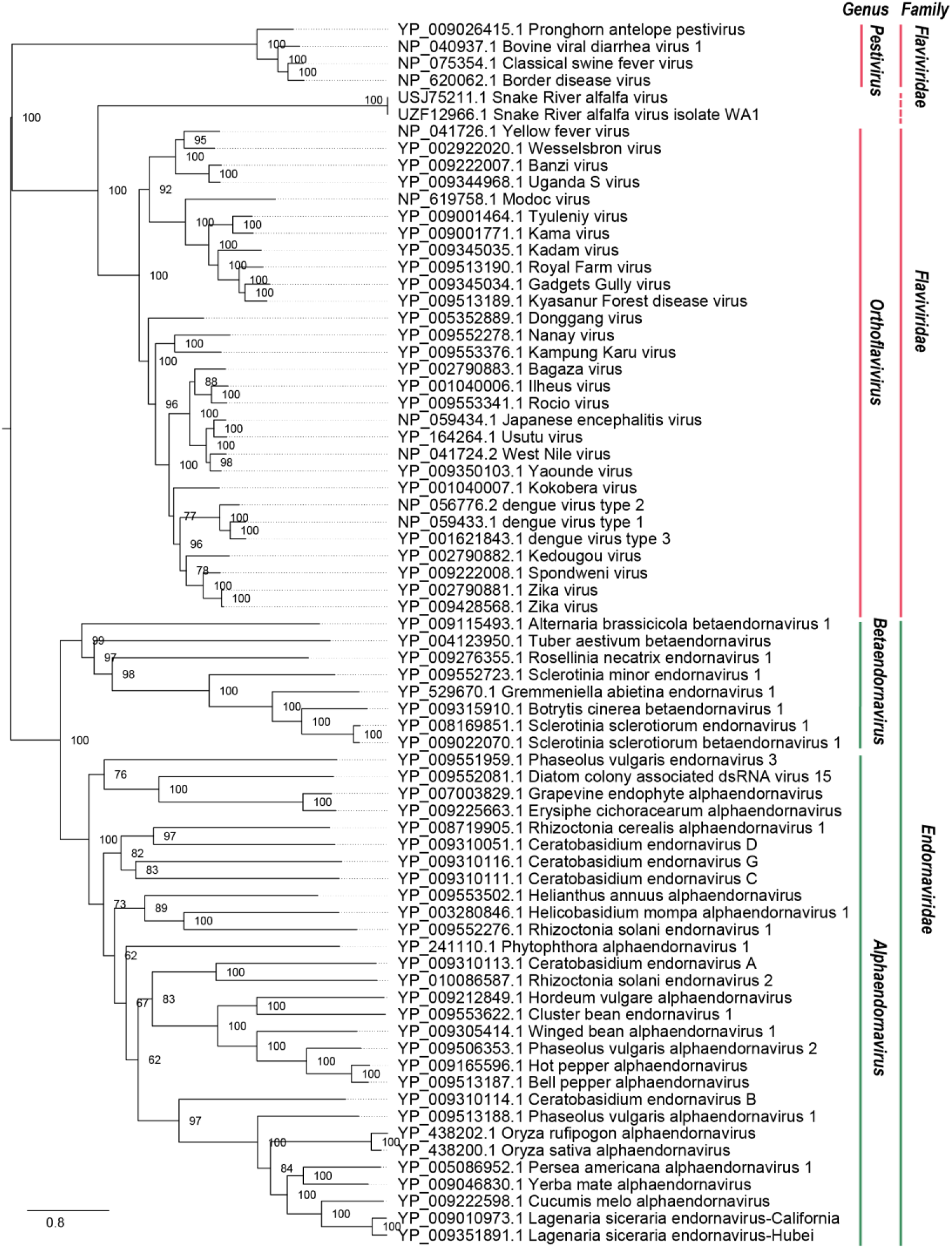
Maximum likelihood (ML) Phylogenetic tree inferred from the same dataset as the article challenging SRAV’s taxonomic status (Postnikova et al. 2023). The clades in red refer to the *Flaviviridae*, and the clades in green refer to family *Endornaviridae*.

**Fig. S4.**
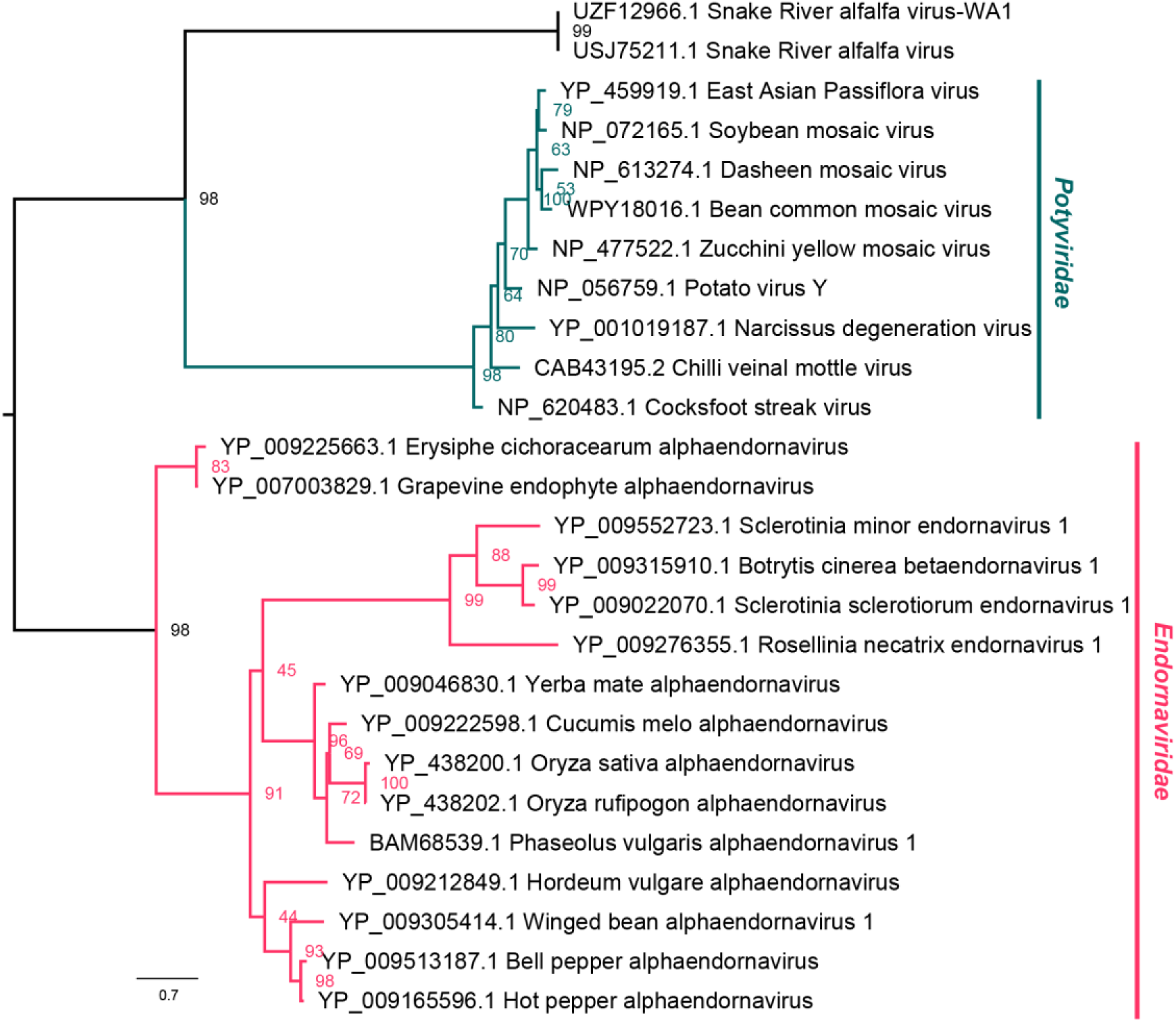
Maximum likelihood (ML) Phylogenetic tree from the RdRP domain of SRAV and representative members of family *Potyviridae* and *Endornaviridae*.

**Fig. S5.**
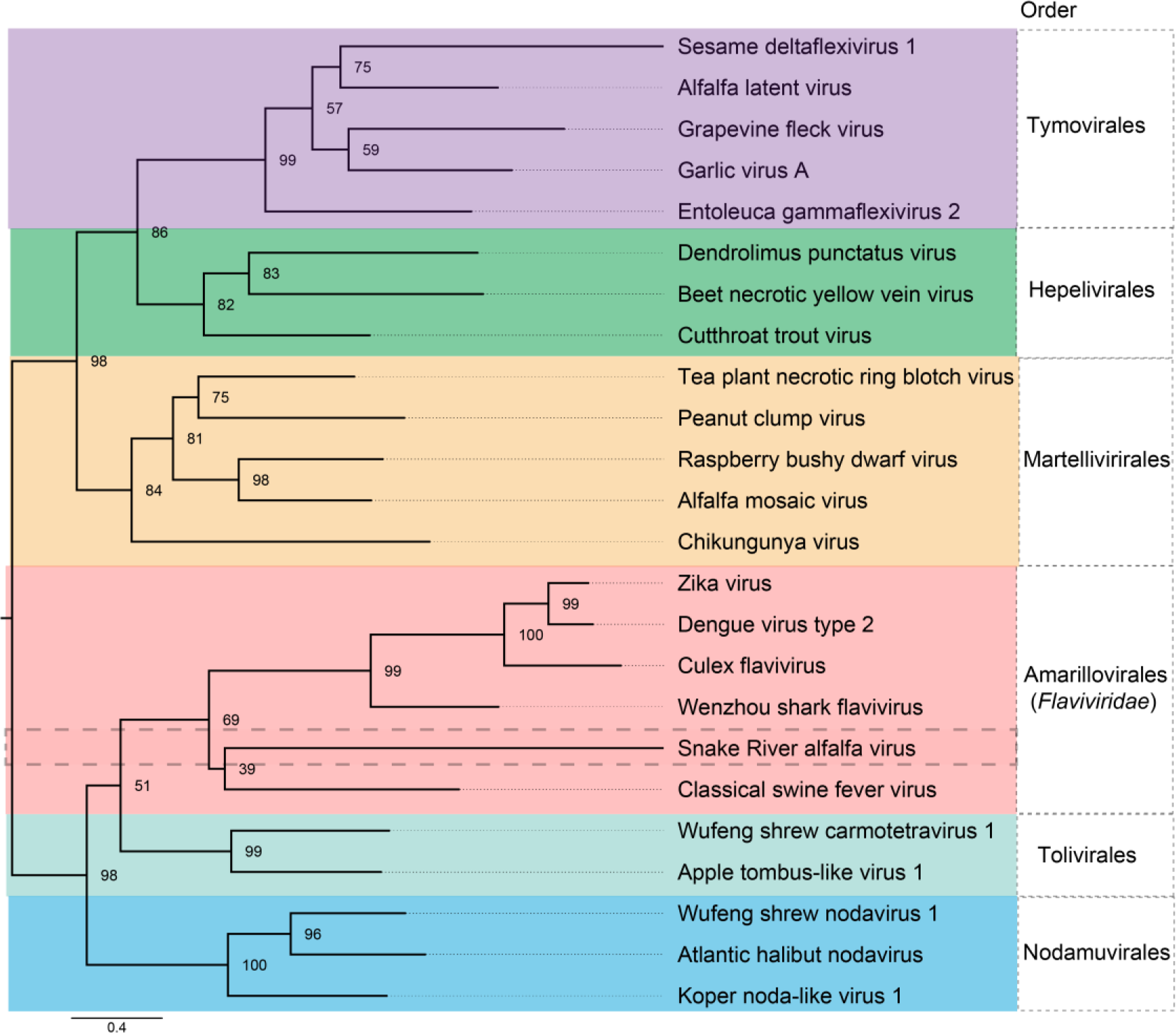
Maximum likelihood (ML) Phylogenetic tree from the RdRP of SRAV and representative members of different order within the phylum *Kitrinoviricota*.

